# Fate-mapping and functional dissection reveal perilous influence of type I interferon signaling in mouse brain aging

**DOI:** 10.1101/2024.05.20.595027

**Authors:** Ethan R. Roy, Sanming Li, Yanyu Wang, Wei Cao

## Abstract

Although aging significantly elevates the risk of developing neurodegenerative diseases, how age-related neuroinflammation preconditions the brain toward pathological progression is ill-understood. To comprehend the scope of type I interferon (IFN-I) activity in the aging brain, we surveyed IFN-I-responsive reporter mice and detected age-dependent signal escalation in multiple brain cell types from various regions. Selective ablation of *Ifnar1* from microglia in aged mice significantly reduced overall brain IFN-I signature, dampened microglial reactivity, lessened neuronal loss, and diminished the accumulation of lipofuscin, a core hallmark of cellular aging in the brain. Overall, our study demonstrates pervasive IFN-I activity during normal mouse brain aging and reveals a pathogenic role played by microglial IFN-I signaling in perpetuating neuroinflammation, neuronal dysfunction, and molecular aggregation. These findings extend the understanding of a principal axis of age-related inflammation in the brain, and provide a rationale to modulate aberrant immune activation to mitigate neurodegenerative process at all stages.

## Main Text

Brain aging is associated with progressive cognitive decline accompanied by structural changes and susceptibility to a list of neurodegenerative diseases that share pathological features of protein aggregation and deposition^1-3^. Inflammation represents a core hallmark of aging, among which IFN-I pathway activation has been implicated with aging in all tissues including the brain^3-6^. To comprehend molecular changes brought by brain aging in a non-biased manner, we performed RNA sequencing analysis on bulk cortical tissues from aged C57BL/6 mice (24 months) compared to young mice (3 months). Not surprisingly, the most significantly upregulated transcripts belonged to inflammatory pathways, such as *C4b, Clec7a, Serpina3n*, and *Cst7* (Extended Data Fig. 1a). Gene Ontology pathway analysis of all differentially expressed genes (DEGs) revealed three networks of pathways predominantly perturbed by aging: one involving immune response (node I), one involving active neuronal signaling pathways (node II), and one involving central nervous system (CNS) development (node III) (Fig. 1a). While pathways in node I were universally enriched in upregulated genes in aged brains, pathways in nodes II and III were mostly enriched in downregulated genes (Extended Data Fig. 1b), suggesting an age-associated uprising of neuroinflammatory response with concomitant depression in pathways relating to neuronal health and physiology. Analysis of Reactome pathways yielded similar results, showing upregulation of inflammatory pathways and downregulation of neuronal and synaptic pathways (Extended Data Fig. 1c).

**Figure 1:**
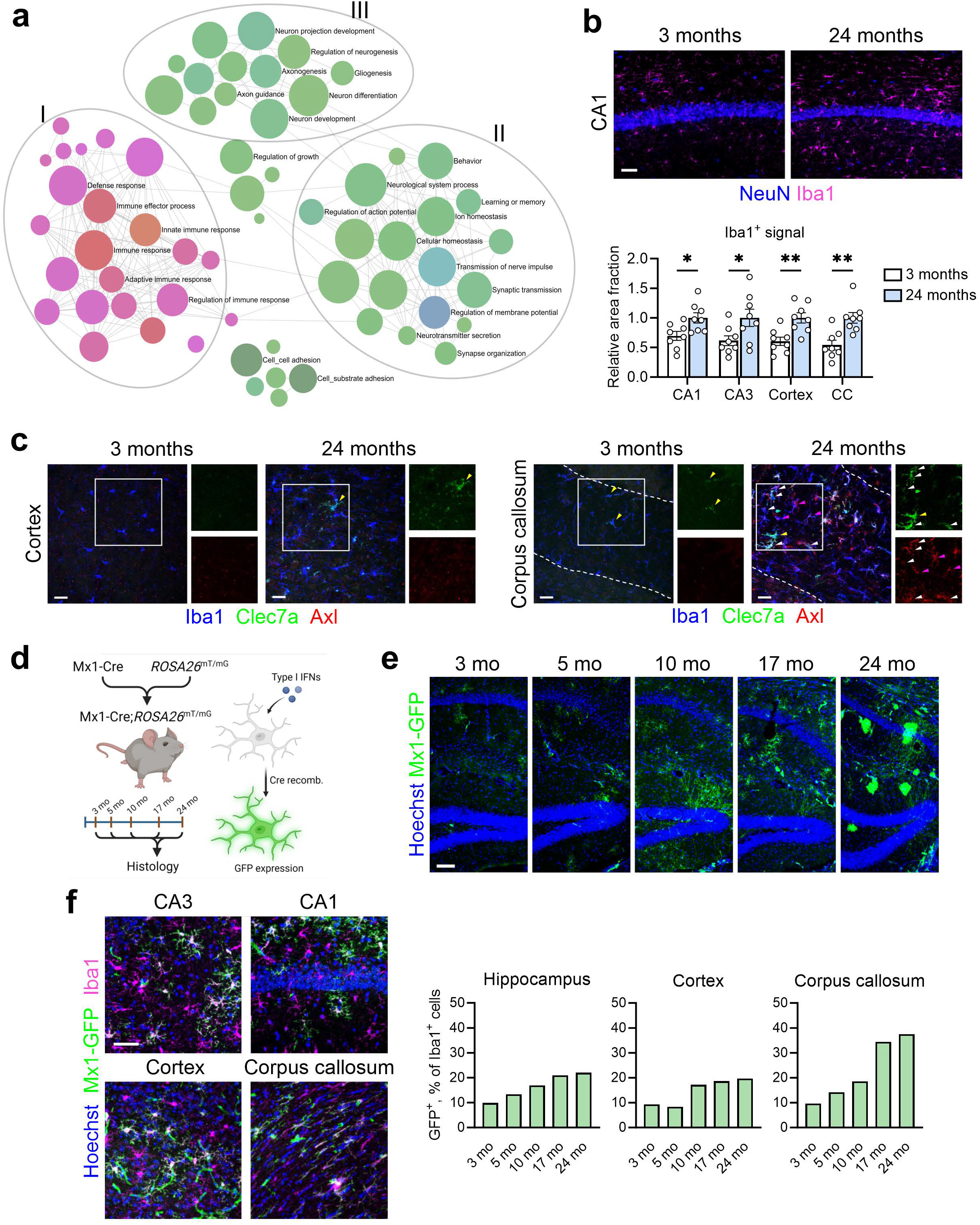
Brain aging entails IFN-I activation in microglia and other cell types, along with gross transcriptomic changes. **a**, Pathway analysis (Gene Ontology) of differentially expressed genes (DEGs) in bulk cortical tissue between aged mice (24 months, *n* = 8) and young mice (3 months, *n* = 10). Selected pathways are labelled. Pink denotes pathways with mostly upregulated DEGs (node I), and green denotes those with mostly downregulated genes (nodes II and III). **b**, Representative images of microglia in CA1 region of young and old mice. Scale bar, 50 µm. Quantification of Iba1^+^ reactivity, expressed as relative area fraction, in various brain regions (3 months, *n* = 8 animals; 24 months, *n* = 8 animals). Data represent means and s.e.m. Statistics were performed with two-tailed *t*-tests. \**P* < 0.05; \*\**P* < 0.01. **c**, Representative images of microglial reactivity markers in cortex (*left*) and corpus callosum (*right*) of young (3 months, *n* = 8) and aged (24 months, *n* = 8) mice revealing multiple subtypes of reactive microglia, such as Iba1^+^Clec7a^+^ (yellow arrowheads), Iba1^+^Axl^+^ (magenta arrowheads), and Iba1^+^Clec7a^+^Axl^+^ cells (white arrowheads). Isolated Clec7a and Axl channels of the same magnification at right. Scale bars, 25 µm. **d**, Schematic showing IFN-inducible reporter model based on spontaneous Mx1-Cre-driven recombination in the *ROSA26*^mT/mG^ reporter line. **e**, Representative images of Mx1-GFP^+^ cells appearing throughout the lifespan in the hippocampus. 3 months, *n* = 4 animals; 5 months, *n* = 4 animals; 10 months, *n* = 2 animals; 17 months, *n* = 4 animals; 24 months, *n* = 12 animals. Scale bar, 100 µm. **f**, Representative images of Mx1-GFP^+^Iba1^+^ microglia in gray matter (hippocampus and cortex) and white matter (corpus callosum) of 24-month-old animals (representative of *n* = 12 animals). Scale bar, 50 µm. Quantification of GFP^+^ percentages of all microglia by age and brain region (hippocampus, *n* = 3,650 cells across ages; cortex, *n* = 3,133 cells across ages; corpus callosum, *n* = 1,641 cells across ages. 3 months, *n* = 4 animals; 5 months, *n* = 4 animals; 10 months, *n* = 2 animals; 17 months, *n* = 4 animals; 24 months, *n* = 12 animals.).

Earlier studies identified prominent IFN-I signature in the choroid plexus as well as microglia isolated from the aging brain^4,7,8^. In bulk cortical tissue, conspicuous upregulation of IFN-I-induced genes (ISGs) was evident (Extended Data Fig. 1a), in keeping with the elevated “innate immune response” and “defense response” pathways (Fig. 1a). To delineate the drivers within the neuroinflammatory landscape in aged brain, we performed transcription factor (TF) motif analysis on DEGs. Remarkably, among the multiple TFs recognizing the motifs enriched in upregulated genes, overrepresented are a host of interferon regulatory factors, including IRF1, IRF2, IRF9, and ICSBP, all of which bind to interferon-stimulated response elements^9^ (Extended Data Fig. 1d). Additional TFs identified included members of the Krüppel-like family (KLF) which may be involved in proliferation, the Zic family, and MAF. Therefore, the parenchyma of the aging brain is molecularly imprinted with rising IFN-I signaling and inflammation.

We next probed age-related, regional phenotypes of microglia, which are known to be key drivers of neuroinflammation^10^. A histological survey detected elevated Iba1^+^ microgliosis in both gray and white matter regions (Fig. 1b,c and Extended Data Fig. 1e). In contrast, Axl and Clec7a, markers of activated microglia associated with Alzheimer’s pathology^11^, were most profoundly expressed by microglia in the white matter, including corpus callosum and fimbria (Fig. 1c and Extended Data Fig. 1e). Upregulation of these markers was heterogeneous, similar to what we observed in amyloidosis brains^12^. This finding of heightened white matter immune reactivity is in agreement with recent reports on glial phenotypes during brain aging^13,14^, which may indicate specialized functions such as clearance of degenerated myelin during aging^15^. This underscores the importance of investigating specific alterations associated with regional aging processes.

Previously, we characterized IFN-I activation in the brains of mouse models of amyloidosis, which is initiated among plaque-associated microglia^12,16^. To assess which cell types are susceptible to such activation during normal brain aging, we generated animals containing an IFN-I-inducible fate-mapping reporter system (Fig. 1d)^12^. Briefly, we crossed mice containing IFN-I-dependent Mx1-Cre with the *ROSA26*^mT/mG^ strain, yielding cells irreversibly labelled with GFP after recombination, and aged these mice to multiple time points. At 3 months, GFP^+^ microglia could be detected in various brain regions, such as the hippocampal layers (Fig. 1e, f). GFP^+^ microglia increased in frequency along the life span in both gray matter areas such as hippocampus and cortex, where they accounted for roughly 20% of all microglia by 24 months, and in white matter areas such as corpus callosum, where they accounted for nearly 40% of all microglia (Fig. 1f).

Other GFP^+^ cell types were also observed in aged animals (Fig. 1e and Extended Data Fig. 2a), such as GFAP^+^GFP^+^ astrocytes which appeared around middle age and were more common by 24 months (Extended Data Fig. 2a); NeuN^+^GFP^+^ neurons, which followed a similar trajectory and were visible in the hippocampal layers, superficial layers of the cortex, and in axon fiber tracts (Extended Data Fig. 2a,b); CC1^+^GFP^+^ oligodendrocytes, which appeared in both aged gray and white matter areas (Extended Data Fig. 2a,c). While rare GFP^+^ blood vessels were found at 3 months, numerous GFP^+^ vasculature was present throughout both gray and white matter areas in aged animals, including large penetrating arteries, arterioles and capillaries (Extended Data Fig. 2a,d). Moreover, at the brain border regions, such as choroid plexus and meningeal layers, GFP^+^Iba1^+^ border-associated macrophages were found (Extended Data Fig. 2e).

Aging stimulates a multitude of molecular changes in cells and enhances the accumulation of senescent cells in tissues and organs^3^. In senescent cells, activation of innate immune cGAS-STING pathway leads to an IFN-I response, which constitutes a late module of the senescence-associated secretory phenotype (SASP)^17,18^. Aging brains are well known to harbor senescent neurons as well as glial cells^19,20^. Recently, microglial-intrinsic cGAS-STING signaling was shown to operate as a driver for neuroinflammation and neurodegeneration in the aging brain^21^. Despite a linear cGAS-STING-IFN-I axis, the functional impact of microglial IFN-I signaling on brain aging has yet to be elucidated. Because it is challenging to silence over a dozen murine genes encoding the type I interferon cytokines, we chose to examine the necessity of the pan-IFN-I receptor (IFNAR), which is essential for a full-scale IFN-I response^22^ and to allow for cell type-specific mechanistic dissection. Hence, we generated 24-month-old mice with conditional deletion of *Ifnar1* in microglia, termed “MKO” (Fig. 2a). Bulk cortical RNA sequencing of MKO animals revealed an overall reversal of the ISG signature seen in aged animals (Fig. 2b), as well as downregulation of pathways related to immune response, indicating an immune-modulatory effect of microglial *Ifnar1* deficiency (Fig. 2c and Extended Data Fig. 3a). TF motifs relating to IRFs were heavily enriched in downregulated genes, as were motifs relating to other aspects of microglial reactivity, such as Spi1 (encoding PU.1) and NFκB (Extended Data Fig. 3b). These data indicate that *Ifnar1*-dependent signaling in microglia contributes most of the IFN-I signature in aged brain tissue, and has broad effects on multiple neuroimmune signaling pathways.

**Figure 2:**
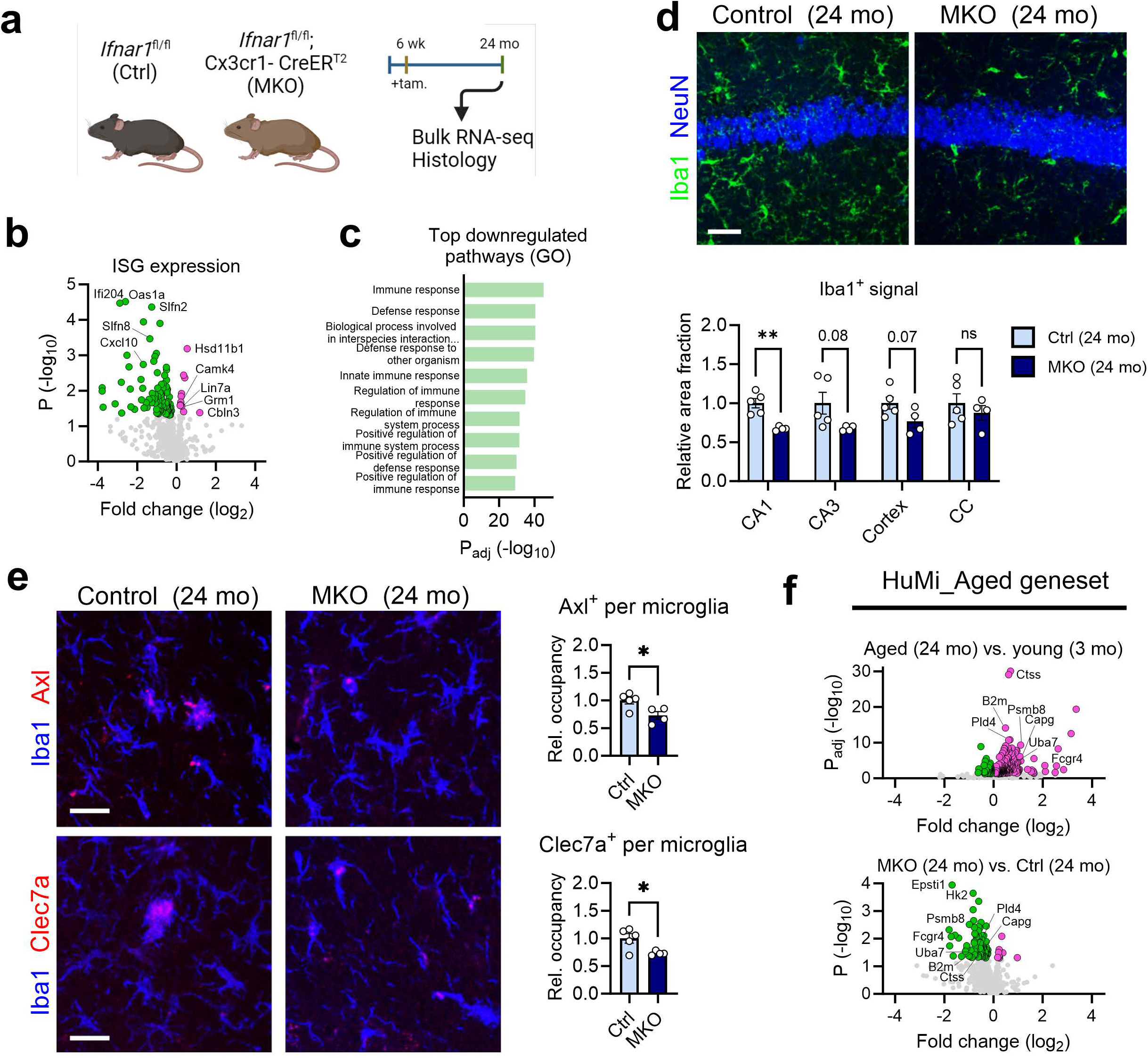
Microglial IFN-I signaling contributes to neuroinflammation of the aging brain. **a**, Schematic showing conditional deletion of *Ifnar1* in microglia through Cx3cr1-CreER^T2^-dependent recombination. **b**, Volcano plot showing expression of a set of interferon-stimulated genes (ISGs) in bulk cortical tissue of aged microglial *Ifnar1* knock-out (MKO) animals (24 months, *n* = 5) compared to aged controls (24 months, *n* = 6). **c**, Top 10 pathways (GO:BP, sorted by significance) enriched in downregulated DEGs (*P*<0.05) in aged MKO brains (24 months, *n* = 5) compared to aged controls (24 months, *n* = 6). **d**, Histological assessment of Iba1^+^ microglial reactivity in control (*n* = 5) and MKO (*n* = 4) brains (CA1 region shown). Scale bar, 25 µm. Quantification of Iba1^+^ area fraction, relative to aged control animals, across various brain regions. Data represent means and s.e.m. Statistics were performed with two-tailed *t*-tests for each region. \**P* < 0.05; \*\**P* < 0.01; \*\*\**P*<0.001; ns, not significant. **e**, Histological assessment of microglial reactivity markers (Axl and Clec7a) in the corpus callosum of aged control (24 months, *n* = 5) and aged MKO (24 months, *n* = 4) brains. Scale bars, 25 µm. Quantification of relative occupancy for each marker within Iba1^+^ cells. Data represent means and s.e.m. Statistics were performed with two-tailed *t*-tests. \**P* < 0.05. **f**, Volcano plots showing expression of a published signature of 978 genes (termed HuMi_Aged) elucidated from aged human microglia ^23^ in (*top*) aged wild-type animals (24 months, *n* = 8) compared to young wild-type animals (3 months, *n* = 10), and (*bottom*) aged MKO animals (24 months, *n* = 5) compared to aged controls (24 months, *n* = 6).

Histological assessment of microglial markers revealed a reduction of Iba1^+^ reactivity in gray matter compartments in MKO brains (Fig. 2d). In white matter, we detected a partial reduction in the amount of Axl^+^ and Clec7a^+^ expression in microglia, amidst comparable Iba1^+^ reactivity (Fig. 2e). Astrocyte reactivity, as measured by GFAP^+^ signal abundance, was not significantly altered in MKO brains (Extended Data Fig. 3c). Finally, we probed our transcriptomic data with a previously published gene list derived from aged human microglia, termed HuMi_Aged^23^, to assess changes in the aging phenotype more generally. Whereas aged wild-type animals displayed a clear upregulation of many of the genes in this module, aged *Ifnar1* MKO animals showed a marked reduction in many of the genes (Fig. 2f), suggesting that microglial IFN-I signaling contributes profoundly to the aged microglia phenotype. A similar effect was observed in genes derived from aged mouse microglia in a separate study^24^ (Extended Data Fig. 3d). Overall, these data support a critical role of microglial IFN signaling in promoting age-related brain inflammation.

Because we observed IFN-I-responsive neurons in aged brain, next we aimed to characterize age-associated neuronal disruption. While neuronal pathway modules were evidently perturbed (Fig. 1a and Extended Data Fig. 1b,c), further analysis showed that downregulated genes in aged brains contained enriched regulatory motifs associated with early growth response (EGR) TFs, which are known to be important in neuronal function, the KROX family, the Krüppel-like family members, MOVO-B, which is involved in neural crest development, and the AP2 family, which are important during neural development (Extended Data Fig. 1d). Besides transcriptomic changes, we detected a significant reduction of hippocampal neuron layer thickness at both CA1 and CA3 regions in aged animals, indicating overt local neurodegeneration (Fig. 3a). Intriguingly, analysis of genes upregulated in *Ifnar1* MKO tissues identified rescued pathways relating to various aspects of neurobiology, including synaptic organization and function, neuronal function, and behavior (Extended Data Fig. 4a,b). CREB (cAMP response element-binding protein) is a key transcriptional regulator of neuronal function, the perturbation of which has been implicated in AD and other CNS disorders^25^. Genes restored in MKO brains were enriched with TF motifs related to the CREB family (CREB, CREM, and ATF-1) (Fig. 3b), and included many target genes of this pathway, including *Bdnf* (Fig. 3c). Another TF family known to be perturbed in the diseased CNS is the MEF2 family. Specifically, Mef2c is a known cognitive resilience factor in AD and controls a network of genes relating to neuronal function^26,27^. Interestingly, *Ifnar1* MKO animals displayed increased expression of both *Mef2c* and *Mef2a*, as well as a number of their target genes (Fig. 3d). Of note, restoration of Mef2c expression by stifling IFN-I signaling mirrors the effect of cGas inhibition in a tauopathy model^28^. Moreover, a list of other known genes relating to neuronal function were also elevated in *Ifnar1* MKO tissues (Extended Data Fig. 4c). In line with these changes, we further found that the layer thickness of CA1 was restored in *Ifnar1* MKO brains compared to aged control brains, with a similar trend in CA3 (Fig. 3e), indicating a reversal of age-dependent neuronal loss, perhaps via decreased microglial removal or de-repression of neurogenesis capacity^4^. In addition, we observed mildly increased density of pre- and post-synaptic puncta in different brain regions of *Ifnar1* MKO animals (Extended Data Fig. 4d), consistent with the findings from the pathway analyses (Extended Data Fig. 4a,b). Overall, our results collectively suggest that microglial interferon signaling is a key contributor to neuronal dysfunction and degeneration in the aging brain.

**Figure 3:**
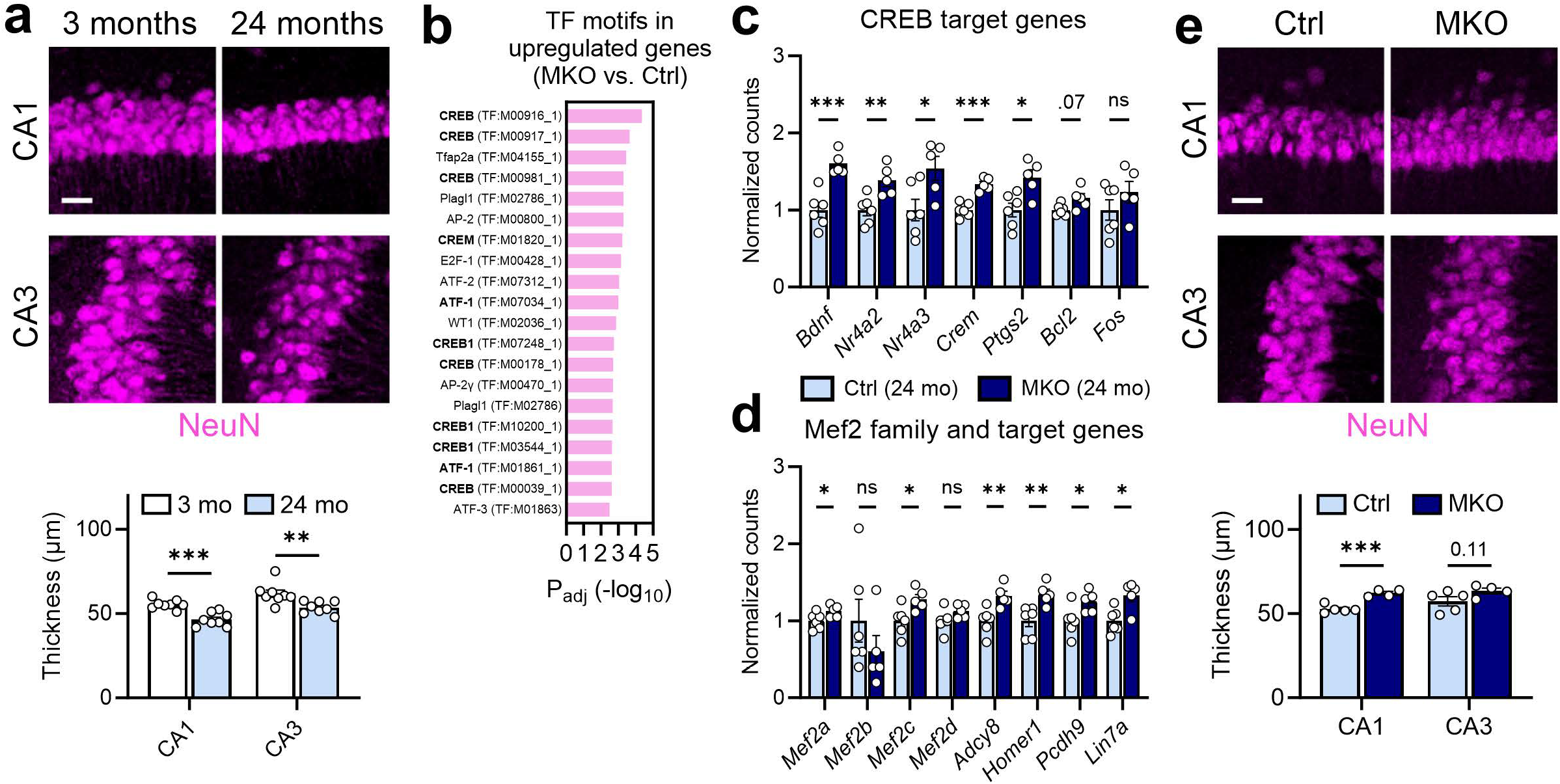
Microglial IFN-I signaling perturbs neuronal function and health during aging. **a**, Histological assessment of NeuN^+^ neuronal nuclear layers in CA1 and CA3 of hippocampus in young (*n* =8) and aged (*n* = 8) brains. Scale bar, 25 µm. Quantification of mean thickness (µm) of CA1 and CA3 layer. Data represent means and s.e.m. Statistics were performed with two-tailed *t*-tests. \*\**P* < 0.01; \*\*\**P*<0.001. **b**, Top 20 TF motifs enriched in upregulated DEGs (*P*<0.05) in aged MKO brains (24 months, *n* = 5) compared to aged controls (24 months, *n* = 6). Terms related to CREB are bolded. **c**, Expression data for target genes of the CREB family of TFs between MKO (24 months, *n* = 5) and aged controls (24 months, *n* = 6). Data represent means of normalized counts and s.e.m. Statistics were performed with multiple *t*-tests (Welch correction). \**P* < 0.05; \*\**P* < 0.01; \*\*\**P*<0.001; ns, not significant. **d**, Expression data for Mef2 family TF genes and their transcriptional target genes between MKO (24 months, *n* = 5) and aged controls (24 months, *n* = 6). Data represent means of normalized counts and s.e.m. Statistics were performed with multiple *t*-tests (Welch correction). \**P* < 0.05; \*\**P* < 0.01; ns, not significant. **e**, Histological assessment of NeuN^+^ neuronal nuclear layers in CA1 and CA3 of hippocampus in control (24 months, *n* = 5) and MKO (24 months, *n* = 4) brains. Scale bar, 25 µm. Quantification of mean thickness (µm) of CA1 and CA3 layer. Data represent means and s.e.m. Statistics were performed with two-tailed *t*-tests. \*\*\**P*<0.001.

Another major hallmark of aging, loss of proteostasis is most notoriously linked to age-related neurodegenerative diseases^3,10^. Aging postmitotic cells frequently accumulate lipofuscin, a nondegradable aggregate of oxidized proteins, lipids and metals^29^. In aged mouse brain, we detected significant lipofuscin build-up in both gray and white matter regions, where these autofluorescent compounds were located inside Iba1^+^ microglia throughout aged brains, and within NeuN^+^ neurons, especially in layer V of the cortex and the CA3 (Fig. 4a, Extended Data Fig. 5a-c). As a pathological hallmark of brain aging, lipofuscin formation is a postulated result of compromised cellular proteostasis^29^. Intriguingly, terms such as “Protein folding”, “Protein folding chaperone”, “ATP-dependent protein folding chaperone”, and “Cellular response to heat stress” appeared as top upregulated pathways by microglial *Ifnar1* ablation (Extended Data Fig. 4a,b). Among the individual DEGs belonging to these processes (Fig. 4b), heat shock protein family members *Dnajb1* (encoding Hsp40) and *Hsph1* (encoding Hsp105) were recently identified as genes universally downregulated in multiple brain regions during aging^14^. Histological analysis showed that microglial deletion of *Ifnar1* reduced overall lipofuscin content in hippocampal microglia (Fig. 4c). Lipofuscin primarily accumulates in the lysosomes; microglial CD68^+^ lysosomes decreased in parallel in MKO brains (Fig. 4c). Furthermore, *Ifnar1* MKO brains accumulated less cytoplasmic lipofuscin within hippocampal neurons (Fig. 4d). These findings uncover a previously unknown function of IFN-I, namely interference of overall protein homeostasis in aging brain.

**Figure 4:**
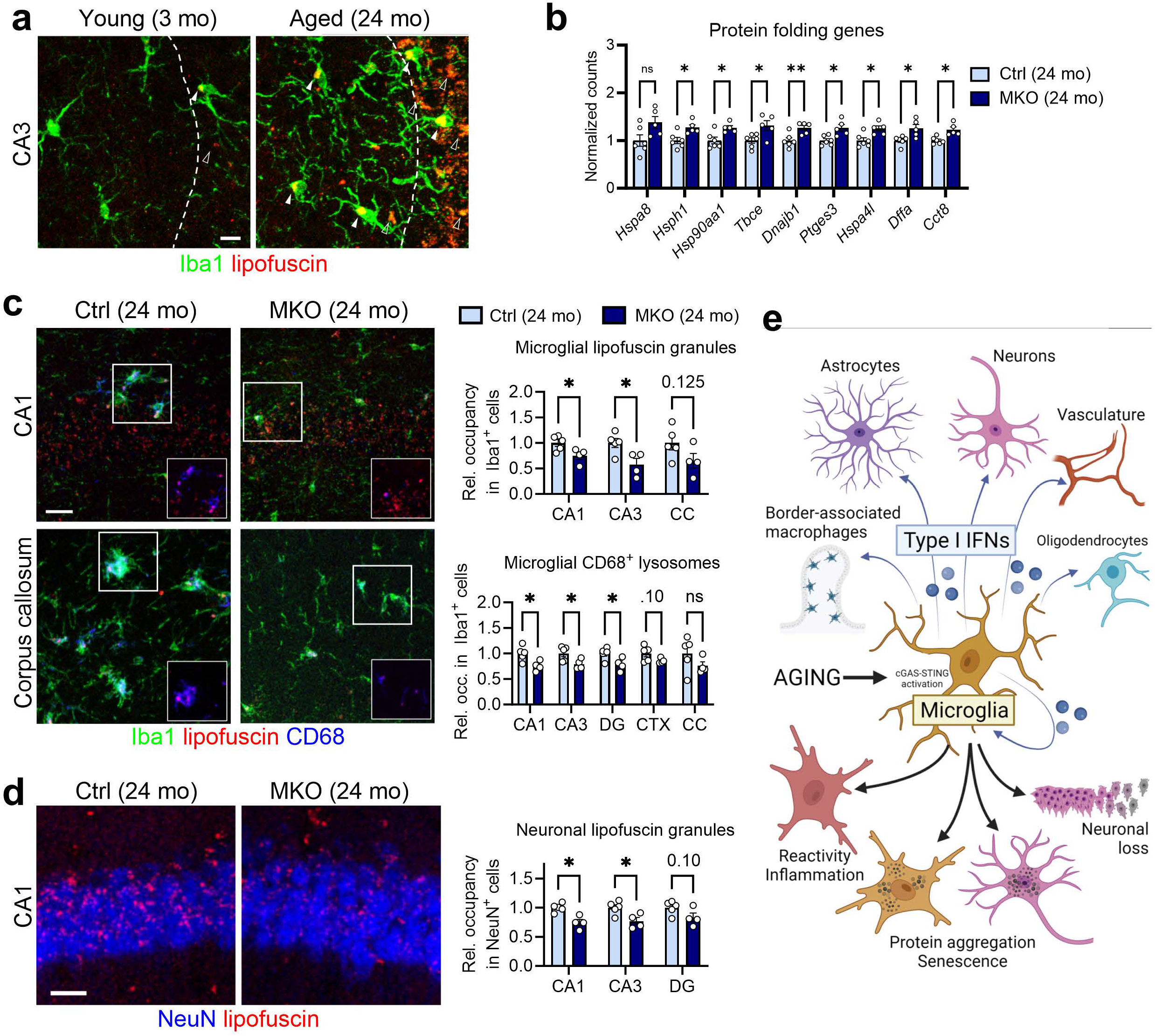
Microglial IFN-I signaling promotes lipofuscin accumulation in aging brain. **a**, Representative images of lipofuscin accumulation in the CA3 region of young (3 months, *n* = 8) and aged (24 months, *n* = 8) brains. Lipofuscin is observed within Iba1^+^ microglia (solid arrowheads) and in other cells (hollow arrowheads). Scale bar, 10 µm. **b**, Expression data for genes related to proteostasis between MKO (24 months, *n* = 5) and aged controls (24 months, *n* = 6). Data represent means of normalized counts and s.e.m. Statistics were performed with multiple *t*-tests (Welch correction). \**P* < 0.05; \*\**P* < 0.01; ns, not significant. **c**, Histological assessment of lipofuscin localized in CD68^+^ vesicles of Iba1^+^ microglia in gray matter (CA1) and white matter (corpus callosum). Scale bar, 25 µm. Quantification of lipofuscin signal occupancy (*top*), and of CD68^+^ vesicle occupancy (*bottom*), in microglia. Ctrl (24 months), *n* = 5 animals; MKO (24 months), *n* = 4 animals. Data represent means and s.e.m. Statistics were performed with two-tailed *t*-tests for each region. \**P* < 0.05; ns, not significant. **d**, Histological assessment of lipofuscin localized in NeuN^+^ neuronal nuclei in the hippocampus (CA1 region shown). Scale bar, 20 µm. Quantification of lipofuscin signal occupancy in neurons. Ctrl (24 months), *n* = 5 animals; MKO (24 months), *n* = 4 animals. Data represent means and s.e.m. Statistics were performed with two-tailed *t*-tests for each region. \**P* < 0.05; ns, not significant. **e**, Graphical representation of aging-induced IFN production, with effects on multiple CNS cell types (*above*), and with specific cellular consequences (*below*), such as neuroinflammation, protein aggregation, and loss of neurons.

Although aging significantly elevates the risk of developing neurodegenerative diseases, the precise role played by age-related neuroinflammation is still incompletely understood. By way of reporter-based fate mapping, we have categorized CNS resident cells of all major lineages in different brain regions responding to IFN-I cytokine during normal aging (Fig. 4e), going beyond previous reports focusing on choroid plexus and microglia. In our survey of 24-month-old brains, IFN-I-responsive microglia accrue up to 20-40% of the total population in different brain areas. As a consequence of senescence downstream of cGas-STING activation, microglial IFN-I response may elicit SASP to fuel tissue wide inflammation. Echoing our findings, other IFN-I-responsive brain cells, such as astrocytes, oligodendrocytes, meningeal immune cells, and endothelial cells, have also been detected under brain aging and disease conditions^12,30-32^. Functional dissection via selective ablation of *Ifnar1* in microglia, in conjunction with histological analyses and transcriptomic profiling, identified microglia as the primary driver of neuroinflammation and IFN-I signature in aging brain, a conclusion in agreement with a recent study^21^. Although TNFα was implicated in neurotoxicity downstream of microglia cGAS-STING activation *in vitro* in that report, our data instead pinpoint an essential role of microglial IFN-I response in promoting neuronal dysfunction and loss in the aging brain. The pathogenic influence of IFN-I on brain network and function in non-aging circumstances has been well documented^33^. Blocking IFN-I signaling within the aged brain was shown to partially restore cognitive function, presumably via reparation of hippocampal neurogenesis^4^. Here, we further identify the CREB and Mef2 families as potential key neuronal transcriptional networks affected by IFN-I in broader areas of aging brain. Inflammation and loss of proteostasis are shared hallmarks of aging and neurodegeneration. Through our examination of *Ifnar1* MKO brains, we discovered an intriguing connection between these two – accumulation of lipofuscin in the brain in part requires full-scale IFN-I signaling, as protein folding machinery is suppressed by chronic IFN-I activation (Fig. 4e). These findings shed new insight on the fundamental forces disrupting the proteostasis network in aging^34^. Senescent cells associated with aging and diseases play a detrimental role in tissue pathology, yet technical limitations hamper definitive identification of these cells histologically or by RNA profiling^35,36^. Nevertheless, lipofuscin has been regarded as one of the markers for senescent cells, especially in the brain^37^. If so, an alternative interpretation of our findings is that IFN-I signaling may further promote senescent cell accumulation (Fig. 4e). Therefore, it is plausible that IFN-I signaling perpetuates a feedforward loop for brain cell senescence, a scenario consistent with recent findings regarding accelerated aging driven by Stat1, the signaling mediator immediately downstream of IFN-I receptor, in progeria mice^38^. As IFN-I response has been pathogenically implicated in neurodegeneration^28,39,40^, this innate immune axis could be a valid target for therapeutic intervention throughout the course of disease development.

## Declarations

### Ethics approval and consent to participate

All animal procedures were performed in accordance with NIH guidelines and with the approval of the Baylor College of Medicine Institutional Animal Care and Use Committee.

### Consent for publication

N/A

### Availability of data and materials

All data generated or analyzed during this study is available from the additional files of this article, or the corresponding author, upon reasonable request.

### Authors’ contributions

ER and WC designed the study, performed the primary interpretation of the data, and wrote the manuscript; ER conducted the experiments and performed the data analysis; SL provided technical support for sample preparation; YW inspected all raw data prior to submission; WC supervised the research and provided funding. All authors contributed and critically reviewed the final version of the manuscript. All authors read and approved the final manuscript.

## Acknowledgements

Support from the BrightFocus Foundation is gratefully acknowledged. We would like to acknowledge technical support from research staff, namely Nadia Aithmitti, Haiying Liu, and Bianca Contreras, and generous sharing of the MXG reporter mice by Drs. Andre Catic and Nicholas E. Propson.

## EXPERIMENTAL MODEL AND SUBJECT DETAILS

### Mice

C57BL/6J mice for both young and aged groups were obtained from the National Institute on Aging (NIA). *Ifnar1*^fl^ (B6(Cg)-Ifnar1tm1.1Ees/J), Cx3Cr1-CreER^T2^ (B6.129P2(Cg)-Cx3cr1tm2.1(cre/ERT2)Litt/WganJ), *ROSA26*^mT/mG^ (B6.129(Cg)-Gt(ROSA)26Sortm4(ACTB-tdTomato,-EGFP)Luo/J), and Mx1-Cre (B6.Cg-Tg(Mx1-cre)1Cgn/J) mice were obtained from Jackson Laboratories. Mice expressing the Mx1-GFP reporter system were generated by crossing *ROSA26*^mT/mG^ reporter mice with Mx1-Cre mice. Mice with conditional deletion of *Ifnar1* were obtained by crossing *Ifnar1*^fl/fl^ mice with Cx3Cr1-CreER^T2^ mice to obtain MKO (*Ifnar1*^fl/fl^;Cx3Cr1-CreER^T2^) and control (*Ifnar1*^fl/fl^) littermates. Mice of both sexes were used for all experiments throughout the study.

For Cre recombination in Cx3Cr1-CreER^T2^-containing mice, tamoxifen was administered to all mice (both Cre and non-Cre) at 6 weeks of age, as described^41^, with modifications. Briefly, 3 mg/ml tamoxifen (Sigma) was dissolved in ethanol and sunflower oil and delivered intraperitoneally to mice (after isopropanol sterilization of the abdomen) daily for five days, with a one-day gap after day 3 to allow animal recovery.

Mice with *ad libitum* access to food and water and were housed in mixed-genotype groups of 3-5 per cage under specific pathogen-free conditions and standard light/dark cycle. Both male and female mice were used in experiments, unless otherwise stated. Mice were analyzed at specific timepoints as noted in the study, and precise ages of all animals used in experiments are listed in respective figure legends. In particular, mice belonging to “aged” groups were all 24 months (±2 weeks) of age. All animal procedures were performed in accordance with NIH guidelines and with the approval of the Baylor College of Medicine Institutional Animal Care and Use Committee.

## METHOD DETAILS

### RNA sequencing

Total RNA from bulk cortical tissues was extracted with Direct-zol RNA Microprep Kits and sent to Novogene Co. (CA, USA) for further processing and bioinformatics. Samples with RNA integrity number >7.0 and 28S:18S ratio >1.8 were used in the analysis. Libraries for sequencing were constructed with the NEB Next Ultra RNA Library Prep Kit (NEB) for Illumina. The library cDNA was subjected to paired-end sequencing with a pair end 125-base pair reading length on an Illumina HiSeq 2500 sequencer (Illumina, San Diego, CA, USA). Gene expression data are included in Supplemental Data 1.

For pathway analysis, the top 500 genes up- or downregulated (based on fold change; *P*_adj_<0.05 for aged vs. young; *P*<0.05 for MKO vs. Ctrl) were queried using the Mouse Molecular Signatures Database (MSigDB). To visualize the relationships between pathways enriched in aging, DEGs (*P*_adj_<0.05) were submitted to the ExpressAnalyst tool (https://www.expressanalyst.ca) and significant Gene Ontology Biological Process (GO:BP) pathways were plotted. Ridgeline plots were also generated using ExpressAnalyst. For TF motif analysis, genes significantly up- or downregulated in aged mice compared to young (all genes with *P*_adj_<0.05), or between MKO and control mice (all genes with *P*<0.05) were queried using the TRANSFAC database in g:Profiler^42^ with significance threshold of 0.05 (Benjamini-Hochberg false discovery rate).

### Immunofluorescence

Mice were perfused with ice-cold saline (0.9% NaCl) after deep anesthesia with ketamine/xylazine, and brains were extracted, fixed overnight at 4°C in 4% paraformaldehyde (Santa Cruz, cat# sc-281692), and dehydrated in 30% sucrose until sectioning. Brains were sectioned into 30-μm tissue sections using a freezing microtome, and tissue was stored in cryoprotectant at −20°C. For staining, floating sections were washed in phosphate-buffered saline (PBS) and blocked for 1 hour at room temperature in a blocking buffer of 10% normal donkey serum (Millipore, cat# S30-100ML) and 1% Triton X-100 in TBS. Primary antibodies were diluted in blocking buffer and applied to sections overnight at 4°C. Tissues were washed with PBS-T (PBS with 0.1% Tween-20) three times, then incubated with fluorescent secondary antibodies diluted in blocking buffer for 1 hour at room temperature. After final washing in PBS-T, sections were mounted on glass slides, allowed to dry, and coverslipped with ProLong Glass Antifade mountant (Life Technologies, cat# P36982). In one experiment, perineuronal nets were visualized with fluorescently labelled Wisteria floribunda agglutinin (Vector Labs, cat# B-1355-2), which was used in conjunction with primary antibody incubation. Tissues from Mx1-GFP reporter animals were stained with anti-GFP antibody to amplify the signal. In certain experiments, slide-mounted tissues from all groups were quenched with TrueBlack autofluorescence quencher (Biotium cat# 23007) after the immunofluorescence procedure to decrease background signals from lipofuscin.

### Image quantification

All imaging in this study was performed with either a Leica SPE or Leica SP8 system. For quantitative analyses, multiple images were used as technical replicates from each animal.

For cell counting in Mx1-GFP animals, multiple images were taken from hippocampus, cortex, and white matter regions of multiple animals per time point. Total Iba1^+^ cells per field, and the portion of them positive for GFP expression, were recorded and combined to estimate % GFP^+^ microglia per region at each age. A total of 8,424 microglia across ages and regions were manually counted for the experiment.

For area fraction calculation, Z-stacks were converted to maximum intensity projections and loaded into ImageJ (NIH) for analysis. ROIs were drawn around relevant areas, images were thresholded manually to overlap fluorescence signals, and percent area was measured. Multiple sections were analyzed as technical replicates for each animal.

For marker co-localization (such as Axl, Clec7a, CD68 and lipofuscin in microglia), Z-stacks (>10 µm total thickness, 1 µm step size) containing cell-type markers (*e*.*g*. Iba1 or NeuN) and markers of interest were generated with confocal microscopy and loaded into Imaris 10.0 software (Bitplane). Using the “co-localization” plug-in, cell-type markers were used as a mask, and the percentage area within the mask of the markers of interest were recorded. Within all co-localization experiments, consistent thresholds were used for each signal to prevent biased measurements.

For assessment of hippocampal layer thickness, confocal images of NeuN^+^ cell layers (CA1 and CA3) were acquired using a 20X lens. For each field of view, the thickness of the layer at three to five points was collected and averaged, and multiple sections were images per animal.

For synaptic density measurements, pre- and post-synapse markers were imaged from the same brain subregions across all animals, due to subtle variations across regions. A 63X oil immersion lens was used to generate Z-stacks (>5 µm total thickness, 0.1 µm step size) which were loaded into Imaris 10.0 software (Bitplane). The “Spots” feature was used to detect synaptic puncta, and the numbers were recorded.

For the CD68/lipofuscin intensity profile, a single-plane image was captured with a 63X oil immersion lens containing Iba1, CD68, and lipofuscin signals. Three channels were split in ImageJ, and intensity along the linear ROI was calculated for each and plotted.

## QUANTIFICATION AND STATISTICAL ANALYSIS

All data in bar plots are presented as means ± s.e.m. Unless otherwise noted, differences between two groups were analyzed by two-tailed Student’s t-tests, and differences between three or more groups were analyzed by one-way ANOVA with Šídák’s post-hoc multiple-comparisons tests, as indicated in figure legends. GraphPad Prism (v10.0.0) was used for all statistical analyses and graph plotting. *P* values less than 0.05 were considered significant (noted as \**P* < 0.05, \*\**P* < 0.01, \*\*\**P* < 0.001 in plots), and those over 0.05 were considered non-significant (“ns”, or numerical P values listed in certain plots). All n values are listed in figure legends for each respective plot. Female and male mice were used for all experiments. All micrographs shown are images representative of multiple replicates as noted in legends. No statistical methods were used to predetermine sample sizes, but similar publications were used as guidelines. Data was assumed to have a normal distribution, though this was not formally determined. Histological experiments were performed and analyzed by a researcher blinded to the genotypes. Biorender.com was used to generate all schematics within the figures.

